# Clustering Protein Binding Pockets and Identifying Potential Drug Interactions: A Novel Ligand-based Featurization Method

**DOI:** 10.1101/2023.05.11.538979

**Authors:** Garrett A. Stevenson, Dan Kirshner, Brian J. Bennion, Yue Yang, Xiaohua Zhang, Adam Zemla, Marisa W. Torres, Aidan Epstein, Derek Jones, Hyojin Kim, W. F. D. Bennett, Sergio E. Wong, Jonathan E. Allen, Felice C. Lightstone

## Abstract

Protein-ligand interactions are essential to drug discovery and drug development efforts. Desirable on-target or multi-target interactions are a first step in finding an effective therapeutic; undesirable off-target interactions are a first step in assessing safety. In this work, we introduce a novel ligand-based featurization and mapping of human protein pockets to identify closely related protein targets, and to project novel drugs into a hybrid protein-ligand feature space to identify their likely protein interactions. Using structure-based template matches from PDB, protein pockets are featurized by the ligands which bind to their best co-complex template matches. The simplicity and interpretability of this approach provides a granular characterization of the human proteome at the protein pocket level instead of the traditional protein-level characterization by family, function, or pathway. We demonstrate the power of this featurization method by clustering a subset of the human proteome and evaluating the predicted cluster associations of over 7,000 compounds.

## Introduction

Whether a drug candidate is targeted at a single protein or multiple proteins, the candidate must also be tested for potential adverse (off-target) effects and toxicity. Targeted assays are the de facto tool to verify the interaction or lack thereof between a new drug with respect to subsets of specific human pathways and proteins.^1^ Using several, narrowly focused assays to assess a new drug’s safety is a response to the complexity of the human proteome. The expanse of proteins coupled with the diversity of their roles and functions is a problem domain too large for drug candidates to be tested *in vitro* against all possibilities. On the computational side, screening large numbers of drug-protein interactions necessitates high-performance computers.^2, 3^ However, even with large amounts of computational power and specialized toxicity assays, unforeseen drug-target interactions and their adverse reactions are frequently detrimental to investigational new drugs (INDs) in clinical trials.^4^

In this work, we demonstrate, using the human proteome, that a pocket characterization-based approach can facilitate the identification of a drug’s most likely targets. We break 4,331 human proteins up into clusters of their pockets using similarities in detected protein-ligand pockets instead of grouping proteins by other characteristics like function or sequence similarity. That is, we focus on evaluating compounds against protein pocket groups – formed by commonalities among the ligands found to bind to those pockets – rather than whole-protein groups typically formed by commonalities among pathway, family, or function. Our method is designed to receive a new compound and identify which groups of protein pockets appear likely to interact with the compound. The output is potentially useful on several fronts, from prioritizing experimental assays to informing *in silico* drug design optimization with regard to potential off-target interactions.

## Background

Describing ligands as keys that match a lock – a protein’s binding pocket – is an oft-used paradigm in computational chemistry.^5^ Drug repurposing, multi-target drugs, and off-target binding analyses are areas that extrapolate from a single ligand and protein pocket to a ligand binding to multiple proteins or different ligands binding to the same protein. ^6, 7^ For multiple ligands to bind to a similar protein pocket, they must share some essential, core configuration of features.^8^ Conversely, for a ligand to bind to multiple protein pockets, the pockets must share some commonality favorable to that ligand. Such cases are often termed “cross-reactivities” or “cross-sensitivities,” where protein pockets which bind similar compounds will in turn have a correlated likelihood of interacting with a similar new drug.^9^ The traditional place to search for cross-reactivities is in similar families, pathways, or proteins with similar functions.^10^ However, in this work we focus on individual pockets, which provide a greater level of detail.

To the best of our knowledge, this approach is the first endeavor in ligand-based clustering of human protein pockets for identification of potential small molecule interactions. This work is inspired by our previous study, ^11^ which demonstrated PDBspheres, a strictly structure-based approach, can help protein function annotation efforts as well as guide in-ference of binding affinity scores from one pocket–ligand pair to another pocket–ligand pair within certain boundaries. Additional related previous work, categorized by methods used for protein- and ligand-based clustering, is summarized below.

### Structure/property-based protein pocket clustering

Analyses based on individual pockets is more granular since a single protein may have multiple pockets. Significant work has been done in this area, especially in clustering protein pockets. Weskamp et al. leveraged shape and physiochemical properties to create a global mapping of cavity (pocket) space and found that similarities in the cavity space are best mapped to ligand binding similarities in comparison with mapping proteins by amino acid sequence or by fold.^12^ Note that the analysis only depends on pocket characteristics; ligand characteristics (binding similarities) are used only for validation. Our approach fuses the ligand and protein pocket feature spaces.

Cavbase, a significant advance in protein pocket clustering, provides a means for comparing pockets based on “pseudocenters,” which are projections of descriptors in 3D space.^13^ Kuhn et al. also applied a principal component analysis (PCA) to their Cavbase similarity matrix for clustering selected MAP kinases. Our approach also applies PCA as a preprocessing step, but to the fused protein pocket and ligand feature space mentioned above.

The CavitySpace database of potential ligand binding sites in the human proteome includes binding site clusters created with a “PMSmax” similarity, which captures shape and chemical similarities between pockets.^14, 15^ The authors iteratively applied the Butina clustering algorithm,^16^ at different PMSmax thresholds. This approach places 31.6% of their database’s 111,330 potential binding sites into one cluster, with the conclusion that “the cavities cannot be classified well.”

It appears that the vast majority of protein pocket featurization and clustering methods rely, as do these studies, on protein features based on pocket structure and/or physicochemical properties^17–21^ – to the exclusion of features based on known interacting ligands.

### Pharmacophore modeling

Pharmacophore modeling includes a spectrum of approaches with applications in drug discovery, lead optimization, target identification, and toxicity prediction^22, 23^ Pharmacophores represent the chemical interactions between a specific ligand or ligands and binding site(s). Pharmacophores can be feature-based^24^ or molecular field-based,^25^ but generally fall into ligand- and structure-based approaches, both of which are related to our approach. For example, protein structure-based methods might infer ideal ligand features from the coordinates of pocket protein residues.^26^ The spirit of this approach is analogous to our aggregation of ligands bound to high-scoring template matches in a pocket, where the similarities of the features in the set of co-complex templates emphasizes the important components of the underlying pocket structure.

In ligand-based pharmacophore models, known active – and sometimes known inactive – compounds are often used to make inferences or sometimes create training data to virtually screen for new inhibitors.^27, 28^ Some ligand-based pharmacophores use clustering on known active and/or inactive compounds to group training sets by similarity.^29^ Our approach also forms clusters using a similarity metric; however, a major difference arises between what is known about the ligands being clustered. Pharmacophores leverage known binders/non-binders for clustering, where our approach clusters known and hypothetical binders based on a pocket’s structure. We cluster protein pockets featurized by ligands from known/predicted template matches. This creates a traversable feature space of protein pocket clusters, while pharmacophore models typically seek to create an in-depth characterization of a particular protein pocket or group of ligands.^30^

### Predicting toxicity, off-target interactions, and adverse drug reactions

A variety of *in silico* approaches exist for toxicity, adverse drug reaction (ADR), and off-target binding prediction; essentially, whether a candidate drug will bind to proteins that are not the drug’s intended target, and what the consequences might be. Methods vary from relation extraction from clinical notes^31^ to hybrid computational pipelines using molecular docking and machine learning.^32^ Pharmacophore modeling is also used, where toxicity and off-target concerns for specific receptors are modeled. ^33, 34^ ADR and toxicity databases like SIDER,^35^ T3DB,^36^ and DrugBank,^37^ have enabled the development and evaluation of numerous machine learning methods. A few notable approaches include REMAP, ^38^ ToxiM,^39^ TargeTox,^40^ and eToxPred.^41^ Recent advances in the toxicity prediction space also include neural fingerprinting,^42^ conformal prediction,^43^ and several others.^44–46^ While our approach doesn’t delve into predicting ADRs, we expect that our simple, granular featurization approach will be useful in multi-target and toxicity prediction.

## Methods

### Pocket Featurization

Our featurization begins with human proteins from the AHA Atlas database^47^ for which a high confidence homology-based structural models can be constructed. For 20,375 proteins from the human proteome (reviewed reference set UniProt^48^ ver. 2020.08.19) the AHA Atlas provides 11,681 structural models created by the homology-based modeling system AS2TS.^49^ The Atlas includes structural models meeting criteria for sequence similarity to and coverage of the reference sequences. Potential binding cavities on these structures are identified by structural matches to ligand binding sites in the whole of PDB^50^ using the PDBspheres library^11^ (Figure 1). While binding sites for individual proteins are sometimes known, many of the structural models in the AHA Atlas do not have solved structures from experiments (i.e., the exact structures or complexes with ligands may not be deposited in the PDB). In this work, we treat pockets identified by PDBspheres as binding sites. Doing so allows for generalization of our approach to any available structural models of a given protein: experimentally solved protein structures, constructed homology models, or structure predictions from methods like AlphaFold. ^51^

**Figure 1:**
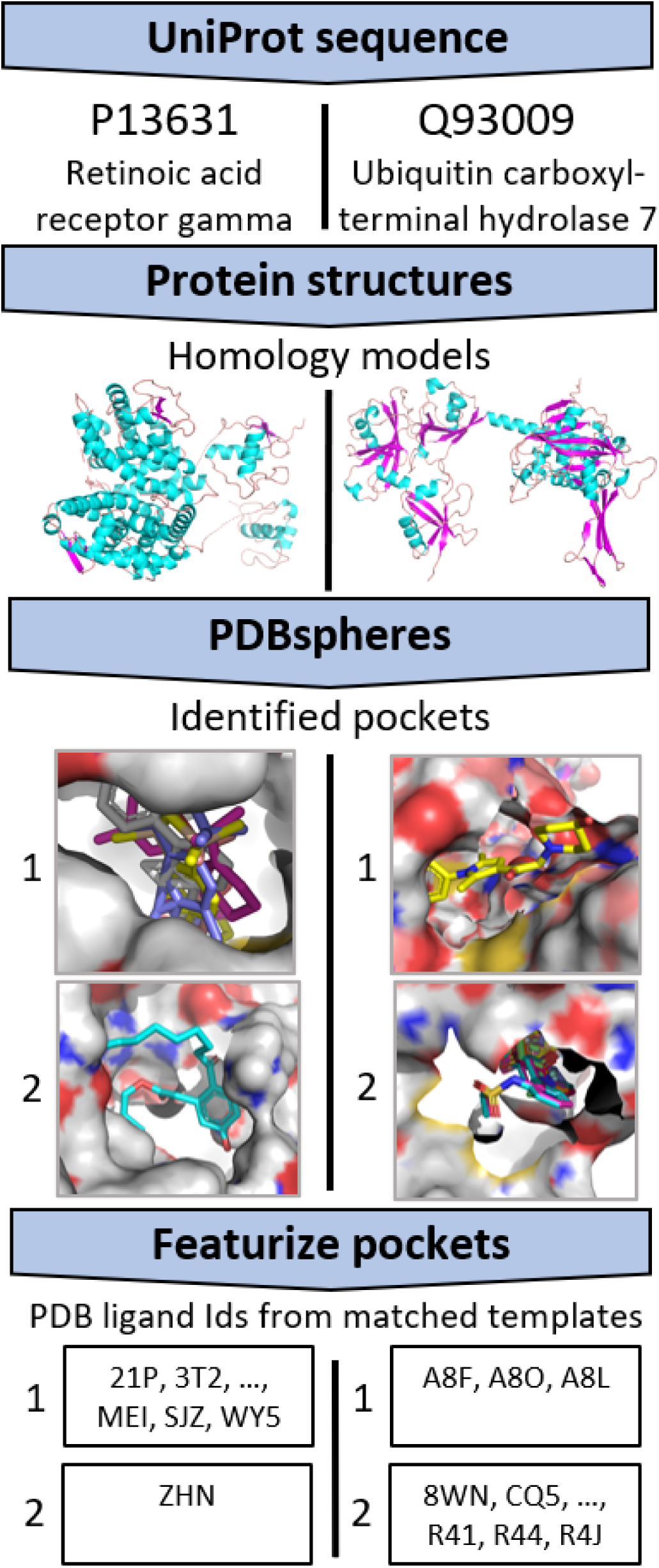
Protein pocket processing and featurization

As mentioned above, the PDBspheres library of binding sites is based on experimentally solved structures of protein-ligand co-complexes extracted from the PDB database. In this work we associate those ligands with the binding pockets identified by protein structural matches between the Atlas human proteome structures and the PDBspheres library entries. This may associate multiple ligands with a particular cavity; the matches depend only on protein-protein comparisons, so an Atlas structure cavity may match multiple cavities from different PDB structures having different ligands. We expect that structurally similar pockets will bind similar ligands, which is a concept explored in our previous work. ^11^ As a corollary, our hypotheses are that (1) ligand similarity provides additional information to characterize pockets (that is, an indication of pocket properties such as charge, hydrophilicity, hydrophobicity, polarity, etc.); and (2) that ligand similarity measures can provide a basis for grouping pockets across proteins. We expect that basing our feature vectors on bound ligands (i.e., not focusing on the pocket structure similarity scores) will avoid possible discrepancies in pocket-ligand clustering which arise from imperfections in the structural conformations of constructed models. Avoiding such pitfalls yields a more reliable and robust model for protein pocket-ligand clustering.

The PDBspheres library^11^ is a comprehensive dataset of all experimentally solved protein-ligand co-complexes that can be extracted from PDB. For this work, we start by excluding from the PDB database (in this case from the PDBspheres library) all entries that may not be relevant to non-covalent binding (principally covalently-bound branched oligosaccharides), may not be biologically relevant (for example, surfactants used in crystallization), or are antibody co-complexes. We filter out non-relevant ligands in three ways. Ligands which overwhelmingly appear as crystallization buffers are outright removed (see supplementary files for ignored antibody structures and crystallization buffers). Covalently-bound oligosaccharides are identified using PDB metadata. Surfactants are identified as ligands that are “surface-bound,” that is, are not substantially within a protein cavity. This is accomplished by constructing a set of spheres tangent to the calculated protein surface^52^ to identify the cavity around the ligand in the PDB biological assembly of the structure used as a basis for the PDBspheres library entry. After removing pockets based on non-relevant ligands, each detected Atlas-protein pocket is associated with one or more ligands; the full set includes 21,948 pockets from the 11,681 proteins – where a single protein might have multiple pockets - and 15,489 different ligands. For an individual protein pocket, we retain a maximum of 20 ligands. In cases where more than 20 template matches are found by PDBspheres, we use a Global Distance Calculation (GDC) to select the best co-complex matches.^53^ Consistent with our hypotheses mentioned above, we use the collection of ligands associated with a pocket as a feature vector or “profile” characterizing that pocket. Figure 1 illustrates different ligand associations for two pockets on two different proteins, Retinoic acid receptor gamma (left) and Ubiquitin carboxyl-terminal hydrolase 7 (right).

As described below, we represent each pocket by a real-valued feature vector derived from the list of ligands associated with the pocket. This serves two purposes. First, we reduce the high dimensionality of both the protein pocket’s structure and properties as well as its associated ligands’ structures and properties to a simple, one-dimensional vector. Second, in characterizing pockets by the small molecules associated with their template matches, our feature space represents a combination of both the ligand binding and pocket structural feature spaces for each entry. Of course, more detailed chemical properties, geometric information, and descriptors may prove beneficial, but we found this foundational method to show significant predictive power on its own.

We assemble our set of pocket-ligand pairs in a 21,948 x 15,489 indicator (binary-valued) matrix, **B**, as Figure 2a shows. This is a sparse matrix where each row represents a human protein pocket and each column is a PDB ligand ID. The columns (15,489 PDB ligand IDs) are provided as a supplemental file. Note that this matrix is predominantly filled with zeros, as 80% of individual pockets are associated with five or fewer PDB ligand IDs.

**Figure 2:**
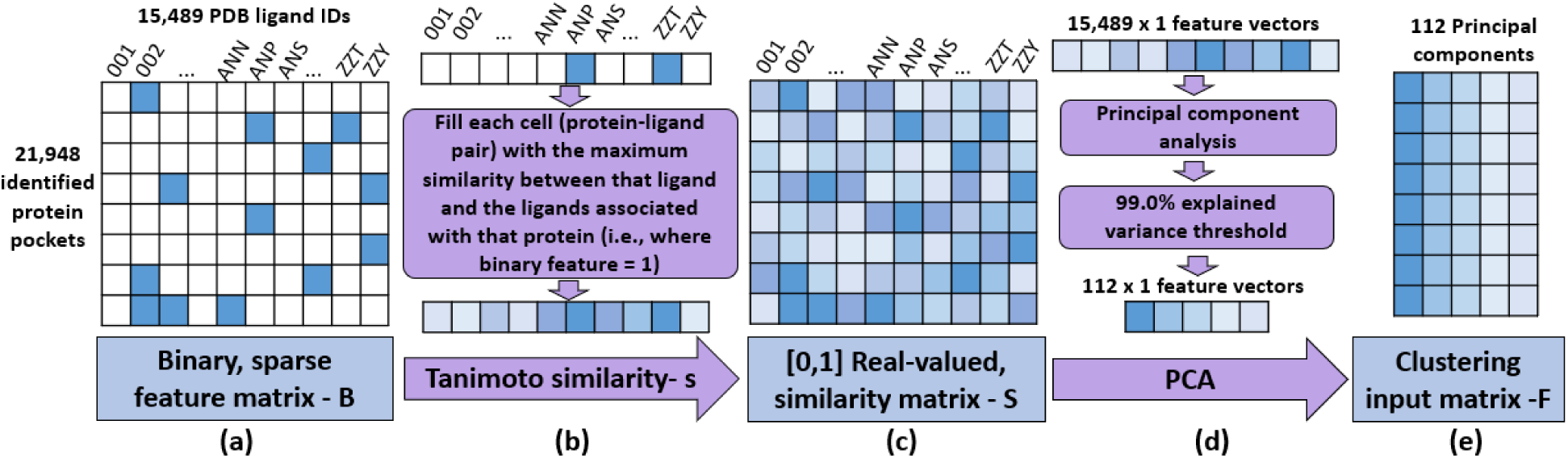
Preparing pocket featurizations for input to clustering

Sparse feature matrices are known to be notoriously difficult for modeling approaches – from logistic regression to deep learning.^54^ Sparse PCA^55^ was the first approach we tried to directly address starting with a binary-valued, sparse matrix. However, due to a lack of orthogonality constraint on the fit components, the scikit-learn^56^ implementation we used was unable to provide explained variance. Therefore, as described in the following paragraphs, we substituted a real-valued similarity measure for the zero-one binary values. The real values are conducive to conventional PCA. The 112 most-significant components of the feature vectors across proteins (that is, across the 21,948 vectors of real values, each 15,489 x 1), explain 99% of the variance of the feature vectors. These reduced-dimension feature vectors are sorted in descending order of explained variance and form the basis for clustering proteins into groups (Figure 2e).

First, we calculate the Tanimoto similarity, which is a [0, 1] continuous value,^57^ between all ligand pairs. Then, for each pocket (row), for each matrix element in that row, we use RDKit^58^ to calculate the maximum similarity between that ligand and all the ligands having a matrix element value of 1 in that row; that is, the maximum similarity between that ligand and any of the ligands that have been associated with that pocket, as Equation 1 describes:

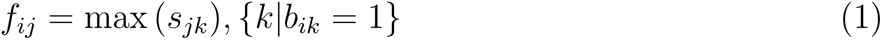

Where:

*f_ij_*: is the real-valued feature [0, 1] for pocket *i*, ligand *j*.

*s_jk_*: is the Tanimoto similarity between ligand *j* and ligand *k*.

*b_ik_*: is the binary-valued indicator whether ligand *k* is associated with pocket *i*.

Equation 1 provides a fast, deterministic computation that introduces more information into the feature matrix, such that every column is no longer an independent feature. This procedure can be thought of as “filling in missing values.” For example, consider the (unrealistic) case where a ligand is duplicated in the set of PDB ligand IDs; that is, the same ligand with different IDs. Assume ligand “*A*” is associated with pocket *i* – either because the protein *i*-ligand “*A*” complex is in PDB, or because a structurally-matching pocket in another PDB protein is in complex with ligand “*A*.” It could be the case that identical ligand “*B*” is not associated with protein *i* because ID “*B*” is not in co-complex with any matching pocket in PDB. In other words, *b_iA_* is 1, and *b_iB_* is 0. Equation 1 sets *T_iB_* = *T_iA_* = 1.0, since the Tanimoto similarity of a ligand to itself is the maximum value, 1.0.

Using the maximum similarity value between a particular ligand *j* and all ligands associated with a particular pocket *i* also accounts for the case where, for example, two very different ligands *m* and *n* nevertheless bind to the same pocket. If ligand *j* is very similar to one or the other of *m* and *n*, then *T_ij_* will be set to the greater of the two similarity values between ligand *i* and ligands *m* and *n*.

Finally, if ligand *j* has very low similarity with any of the ligands associated with pocket *i*, then *T_ij_* will be given a low similarity value (albeit the greatest of its low similarities with the associated ligands). This could be a “false negative;” it could be the case that no ligand similar to ligand *j* has ever been seen in a PDB co-complex having a pocket matching pocket *i*, but nevertheless such a ligand would bind to pocket *i*. The working assumption of PDBspheres is that the set of pockets identified throughout PDB by proximity to a ligand covers – at least with a high degree of similarity – the full universe of protein binding pockets. Each pocket has 15,489 (now real-valued) features, that is, the similarities calculated using Equation 1 yield the similarity matrix **S** (Figure 2c). We implement feature reduction by performing PCA on the feature matrix. This provides both the ability to make an interpretable reduction in dimensionality, and quantifies the explained variance associated with each component produced. While exploring the linear combinations of singular value decompositions that make up each component is outside the scope of this work, using PCA maintains the possibility of doing so.

While the five most-significant components explain 95% of the variance among the original columns, we use the 112 most-significant components; these explain 99% of the variance. We substitute the elements of these components for the original 15,489 features for each pocket. The resulting matrix is suitable for clustering.

### Pocket Clustering

Since we do not have pre-existing estimates of the number of groups of pockets that will best characterize the human proteome – or the subset of the human proteome in the AHA Atlas – we use clustering methods that do not need a target number (prior) of groups to produce. Density-based spatial clustering of applications with noise (DBSCAN^59^) is a clustering algorithm that does not need a number of clusters prior and does not constrain clusters to a specific size. DBSCAN uses the concept of core samples (dense areas) and a distance threshold to form clusters from core samples and nearby non-core samples. The minimum number of points forming a core sample defines how densely populated a neighborhood must be and is a reflection of noise in the dataset. A distance threshold – the maximum distance between cluster members and the core-samples area – controls the size of clusters, therefore influencing the number of clusters and how many points cannot be clustered within the minimum-size constraint; such points are labeled “outliers” (rather than, say “monad clusters”). The simplicity of DBSCAN is attractive for interpretability. DBSCAN retains the concept of Euclidean distance between clusters and points which is captured by the eigenvalues of the principal components. Therefore, the output of a DBSCAN clustering on our feature matrix of components can be analyzed in terms of 112-dimensional Euclidean distance in feature space. It is important to note that by thresholding at 99% of explained variance, examining the distance between points in our feature matrix is an approximation and not numerically exact.

Our goal in choosing values for these hyperparameters was to strike a balance between fewer large clusters – where some very large clusters may be dominant – and more small clusters – creating an unnecessarily large number of small clusters to review. We set the minimum number of pockets necessary to create a cluster in a local neighborhood of feature space to 10. This helps ensure that clusters are sufficiently populated for examination and evaluation of their members’ characteristics. The other major DBSCAN parameter, the core-to-member maximum distance, has significant impact on the DBSCAN output; we experimented with a range of values for the distance threshold, examining the results in terms of the number of clusters, number of outliers, and largest cluster size, as Table 1 reports. Between large and small distance thresholds, DBSCAN marks pockets as outliers when they are far from other pockets or in a feature-space region without the minimum 10 nearby pockets necessary to form a cluster. It should be noted that the DBSCAN algorithm allows the 10-member minimum constraint to be violated in cases where the core sample is sufficiently dense.

**Table 1:**
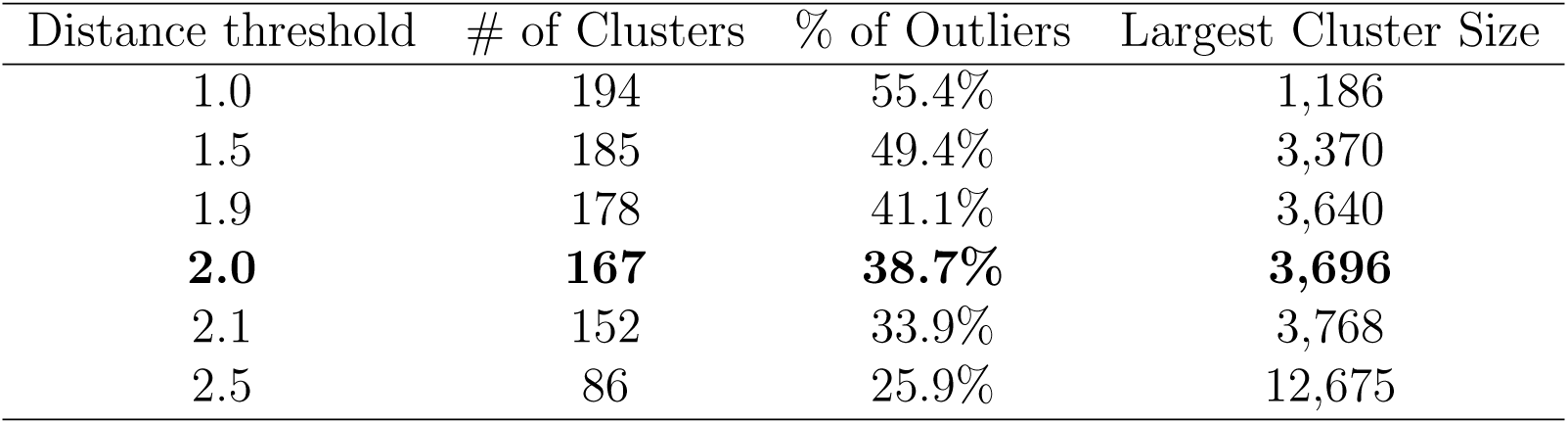
DBSCAN cluster results on 21,948 protein pockets with varying maximum core-to-member distance threshold

| Distance threshold | # of Clusters | % of Outliers | Largest Cluster Size |
| --- | --- | --- | --- |
| 1.0 | 194 | 55.4% | 1,186 |
| 1.5 | 185 | 49.4% | 3,370 |
| 1.9 | 178 | 41.1% | 3,640 |
| **2.0** | **167** | **38.7%** | **3,696** |
| 2.1 | 152 | 33.9% | 3,768 |
| 2.5 | 86 | 25.9% | 12,675 |

Table 1 displays the clustering results achieved by varying the distance threshold from 1 to 3, which spans the extremes of many clusters with many outliers to few clusters with few outliers. A plateau seems to appear in the 1.9 – 2.1 range where most pockets are in-distribution, the largest cluster sizes are around 4,000, and the number of clusters is relatively steady at approximately 160. While the percentage of outliers is larger than desired, the 10-member minimum constraint and the fact that the data are from a subsample of the human proteome both contribute to pockets appearing anomalous. Focusing on the core functionality of the approach and possibility of adding data in the future, a distance threshold of 2.0 was selected as a reasonable balance among the various factors considered.

## Results

The DBSCAN results with a distance threshold of 2.0 yield 167 clusters of 13,455 protein pockets; the remaining 8,493 pockets are outliers. The first three principal components of the feature space capture 93.4% of the variance across pockets. Figure 3 visualizes these three components spatially and the size of each cluster is illustrated by the size of its scatter point. The 167 clusters vary in size from seven to 3,696 pockets and as mentioned above, the 10-pocket lower bound discussed can be violated in dense regions. The population of 167 clusters has a median of 16 and a mean of 81 pockets per cluster; a right-skewed distribution. As an initial check of the clustering, we examined the clusters in terms of the unique proteins (UniProt IDs) in each cluster. The statistics for unique proteins are similar to those for pockets; the minimum number of unique proteins in a cluster is six and the maximum number of unique proteins in a cluster is 1,768. This indicates that even small clusters group different proteins, thus small clusters are not simply repeated instances of the same pocket from a single protein biological assembly (as in a homodimer, for example). On average, 52 different proteins make up a cluster, where the median is 14 unique proteins, giving a skewness to the population of pockets.

**Figure 3:**
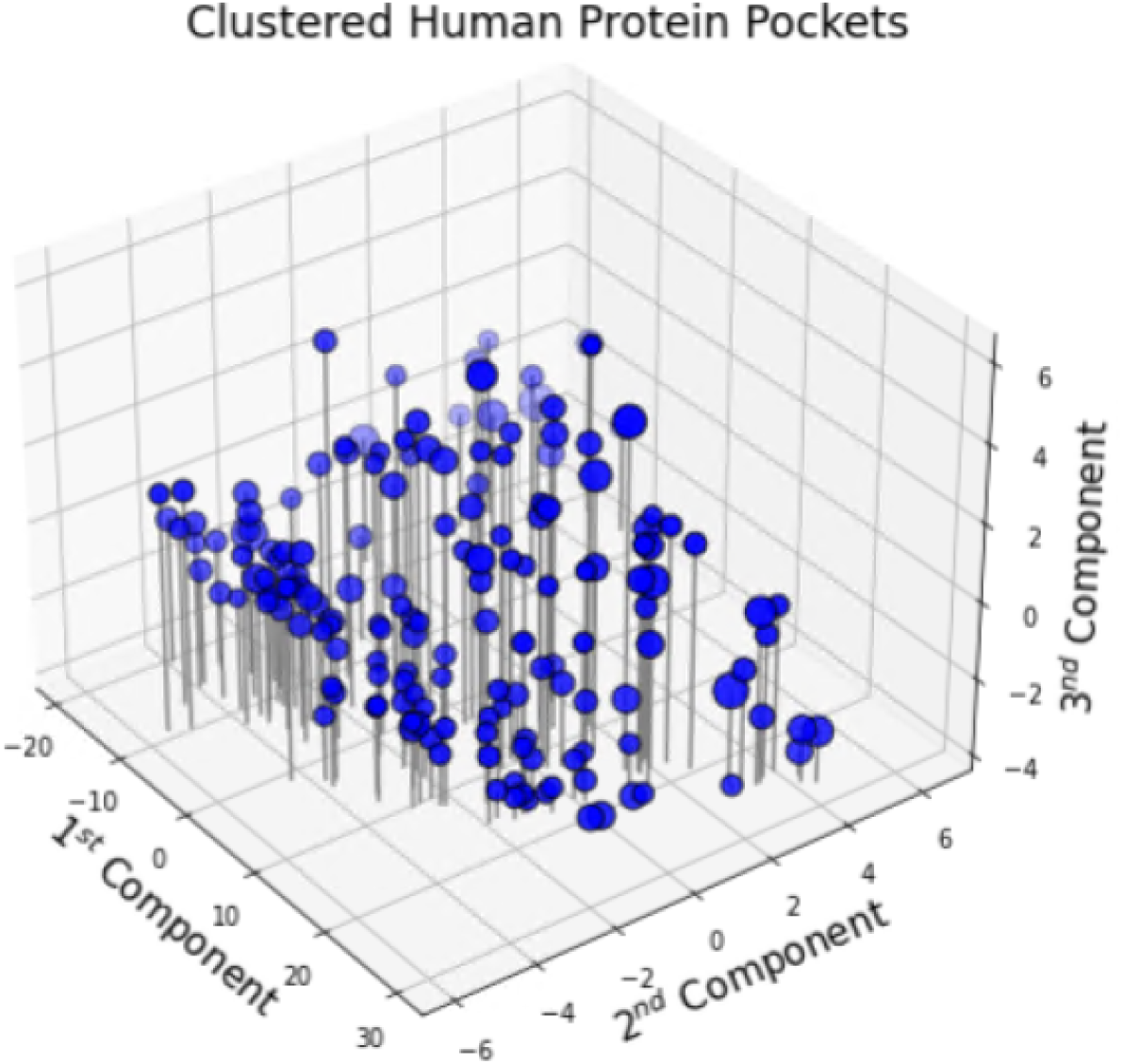
Protein pocket cluster centers (core samples centroids); 167 clusters visualized by their first three principal components and size.

In accord with previous protein-based clustering efforts,^11^ we note several indications that the clusters form biologically meaningful groups. First, we look at cluster members’ Enzyme Commission (EC) numbers,^60^ which classify enzymes by the type of reaction they catalyze. Of the 167 clusters, 54 (32%) consist of proteins having the same major EC class. While typically not all proteins in a cluster have an EC number, nevertheless a significant number of proteins share the same top-level EC number in a significant number of clusters. Furthermore, 36 of the 54 single-major-number clusters consist of proteins also sharing EC subclass and sub-subclass numbers, despite the fact that there is no cluster having fewer than six proteins.

Second, we look at cluster members’ appearance in the signaling pathways identified by the Small Molecule Pathway Database (SMPDB^61^). There are 17 clusters in which all members are associated with proteins in a single pathway; seven of the 17 clusters have more than one protein associated with that same pathway. The pathways associated with those seven clusters are: Ubiquitin-Proteasome, Rac 1 Cell Motility Signaling, and the GnRH Signaling Pathway. Notably, five of these seven clusters are associated with the Ubiquitin-Proteasome pathway which consists of 28 different proteins. In SMPDB, a single protein is often associated with multiple pathways, which makes looking for homogeneous cluster-pathway associations difficult. Filtering clusters for those with multiple proteins associated with the same set of pathways yields 12 clusters of interest. Cluster #29 is one such example; it has 11 pockets from 10 different proteins. The proteins in cluster #29 found in SMPDB are each associated with the same three pathways: Folate Metabolism, Methotrexate Action, and Methylenetetrahydrofolate Reductase Deficiency. This association makes sense as the leading PDB ligand IDs associated with the pockets in cluster #29 are folic acid and methotrexate. While observations such as these based on UniProt IDs, EC number, and SMPDB associations are interesting, a more quantitative validation of the clusters can be achieved.

### DrugBank

The DrugBank database^37^ lists protein-ligand pairs known to interact (that is, bind). Our clustered protein set includes proteins for which DrugBank shows interactions with 4,827 compounds. Of these compounds, those which have known activity (among DrugBank’s categories target, transporter, enzyme, and carrier) with a protein that was successfully clustered. 2,232 exist in our feature set (matrix **B**); that is, our approach used these compounds as protein pocket features. The remaining 2,595 compounds are “novel” to our method.

We look first at the DrugBank compounds found in our feature set. Since these compounds are part of the “training” set (e.g., used in our featurization for unsupervised clustering), they do not provide a basis for unbiased tests of the clustering. Certainly, this is the case if the DrugBank interacting protein-ligand pair appears in PDB as a co-complex.

In cases where that particular protein-ligand pair does not exist in PDB then it is included in our feature set as a result of pocket similarities. Therefore the “training” compounds can help test our approach. We do so by treating these “training” compounds as novel ligands, and determining which protein clusters these ligands would be associated with. Our procedure is this: (1) measure each DrugBank ligand’s similarity with the 15,489 ligands in our feature vector; (2) apply the pre-fit PCA to render 112 x 1 feature vectors used for clustering;. (3) calculate the feature-space distance between each ligand and every DBSCAN core sample; and (4) label each DrugBank compound with the number of that nearest core sample’s cluster. This creates a projection of each compound in the protein pocket feature space and allows for creating an ordered list of the protein pocket clusters nearest each compound.

The first row of Table 2 shows the results for this “training” group of ligands. Of the 2,232 DrugBank compounds that are in our feature set, 1,129 (51%) have a known interaction with a protein in one or more of the nearest 10 clusters from the field of 167 clusters. Further, 726 of those (33%) have all known interactions within their 10 nearest clusters. These results are positive and demonstrate significant predictive power in a relatively simple method across a broad spectrum of drug-like compounds and human proteins. Narrowing down to the five nearest pocket clusters, 44% of the compounds have a known interaction in one or more of the nearest five clusters and 28% have all known targets within those five nearest clusters. Finally, 456 of the 2,232 ligands in this DrugBank subset (20%) are nearest to a cluster which contains all their known interactions. While these are not validations of the exact pocket from a known interaction, which typically is not known, the presence of a protein known to interact with a ligand in a cluster nearest that ligand is preliminary evidence that our approach is functioning properly.

**Table 2:** DrugBank compound results for known interactions in nearby clusters.

| DrugBank |  | One or more interactions in: |  |  | All known activity in: |  |  |
| --- | --- | --- | --- | --- | --- | --- | --- |
| Set | Count | 10 nearest | 5 nearest | Nearest | 10 nearest | 5 nearest | Nearest |
| “Train” | 2,232 | 1,129(51%) | 983(44%) | 727(33%) | 726(33%) | 617(28%) | 456(20%) |
| Test | 2,595 | 839(32%) | 763(29%) | 642(25%) | 529(20%) | 423(16%) | 294(11%) |

Note that there are several reasons why a ligand may not be projected near its known interacting proteins. First, PDBspheres may not have identified a pocket, meaning it is absent from the dataset in the first place. Second, even for all identified pockets, their predicted template matches may be inaccurate. Third, even if a pocket is identified and well-characterized by the predicted templates, it can be poorly captured by the PCA or, fourth, thrown out as an outlier by the clustering method. Finally, ambiguity from the simple featurization of that pocket may also lead to errors in ligand-to-pocket associations. These possible failure points are important to the context of results, but also point to this approach improving as new PDB entries can substitute for homology models and allow additional human proteins to be included in the dataset.

The remaining 2,595 compounds of the DrugBank subset are not part of our feature set, and therefore are not associated with any pocket among our proteins’ clusters; thus, they are “novel” to our method. We call these compounds the “test” set.

The second line of Table 2 shows the results of applying to the “test” set the same procedure as that applied to the “train” set compounds. Of these 2,595 compounds, 839 (32%) are placed with a known target in one or more of the nearest 10 clusters and 20% have all known targets within the 10 nearest clusters, among our global set of 167 possible clusters. For the five nearest pockets, 763 (29%) of the compounds are placed with one or more known targets, and 16% include all known targets. A final point of comparison between the two sets is that 294 drugs are placed in a cluster which includes all their known targets. These percentages are lower across the board in comparison with the “training” set ligands in DrugBank, which is to be expected. However, this set of compounds is a fair test of the generalization of our method.

### Drug Repurposing Hub

The DrugBank compounds provide a macro-level confirmation of the featurization and clusters. Micro-level analysis is possible with the Drug Repurposing Hub database.^62^ Of 6,550 Drug Repurposing Hub compounds that are in the AHA Atlas, the Drug Repurposing Hub indicates there are 2,899 compounds that have a known interaction with a protein present in one of our clusters, but are not included in the ligand feature matrix, **B**. Thus, these compounds are “novel” to our method. Using the criteria from the rightmost columns in Table 2, 405 (14%) have all known interactions within the ten nearest clusters, 334 (12%) are within the nearest five clusters, and 255 (9%) have all known interactions contained by their nearest cluster. The AHA Atlas database provides machine learning (Coherent Fusion^2^), molecular docking (AutoDock Vina^63^), and MM/GBSA scores for many combinations of protein and ligand.^3, 64^ Information from these scoring functions allows for a deeper understanding of which ligands and pockets are well-represented and why. Among the results from the 9% of compounds with any known activity exclusively in their nearest cluster, we filtered the compounds to those predicted in the smallest clusters to extract a list of five examples.

In the first example, two preclinical compounds^1^ (Figure 4, molecules A and B) are present in PDB as co-complexes which match the same protein pocket (in the structural model of UniProt ID Q93009) and were assigned to cluster #58. These compounds are both ubiquitin-specific peptidases. Cluster #58 consists of 67 pockets from 52 different proteins; the most common PDB ligand IDs that featurize these pockets are N5S, N6J, HBI, N5D, and A8O. The pocket that is included in cluster #58 that PDBspheres matched to the pocket of the structural model for UniProt human proteome reference sequence Q93009 has three PDBspheres-predicted ligands which featurize it: PDB ligand IDs A8O, A8L, and A8F^2^. Interestingly, molecules A^65^ and B^66^ (Figure 4) do not have high similarity to any of the 15,489 compounds in our dataset; the dataset compound having the greatest similarity to these two is PDB ligand ID D3U (“2-PCPA derivative”), which has Tanimoto similarities of 0.41 and 0.43 to each, respectively. Cluster #58’s pockets represent the most likely candidates with which these two preclinical drugs have interactions. The protein pockets in that cluster that are most similar to the “spheres” (PDB co-complexes) from which molecules A and B were derived are structural models of UniProt IDs (1) Q93008 (probable ubiquitin carboxyl-terminal hydrolase FAF-X, (2) O00507 (probable ubiquitin carboxyl-terminal hydrolase FAF-Y), and (3) Q9UPU5 (ubiquitin carboxyl-terminal hydrolase 24). Among the receptors in cluster #58, our method highlights these three ubiquitin carboxyl-terminal hydrolases as the most likely proteins to interact with molecules A and B. This information is more specific than finding similar proteins through protein family associations. The proteins of the co-complex pocket (Q93009) and all three potential interacting pockets (Q93008, O00507, Q9UPU5) belong to protein family (peptidase C19) and have the same EC classification (3.4.19.12). The peptidase C19 family^67^ contains 133 different human proteins and there are 782 proteins under the ubiqitinyl hydrolase 1 EC serial number (12), of the omega peptidases sub-subclass (19), in the peptidases subclass, of the hydrolases EC class (3).^68^ Instead of generally associating hundreds of proteins with the known target receptor, our approach provides quantified, granular information about similar receptors without limiting results to ubiquitin carboxyl-terminal hydrolases.

**Figure 4:**
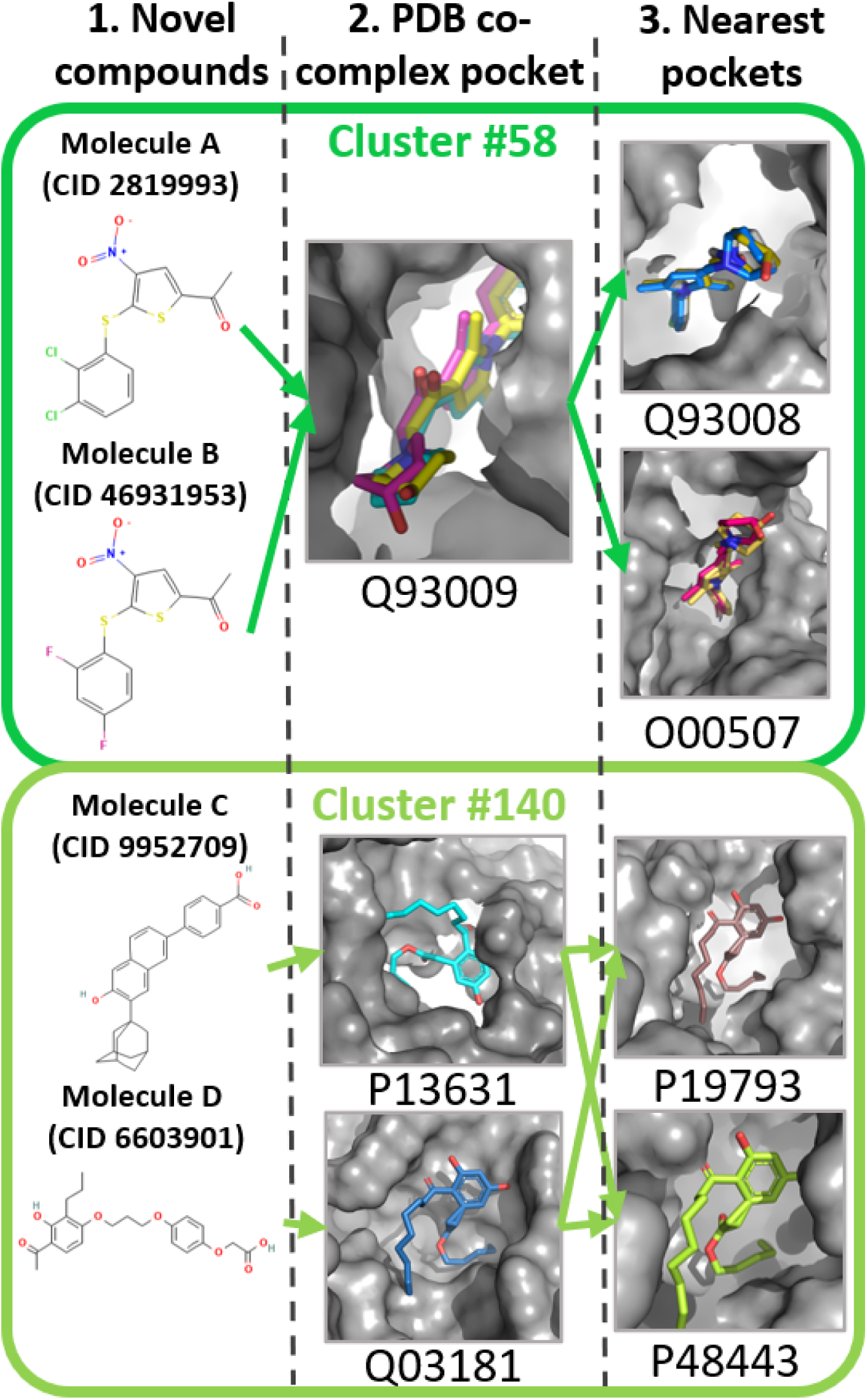
Sample compounds (CID: PubChem compound identifier) that are not in clustering feature matrix B (panel 1); their pose in the protein structural model of the UniProt reference sequence included in clusters #58 and #140 – the poses are derived by alignment of the PDB co-complex pockets that matched the structural model to the structural model of the UniProt sequence (panel 2); and the nearest – in feature space – ligands/structural models, again in poses derived by aligning the PDB co-complex to the structural model alignment (panel 3).

Another pair of preclinical compounds^3^ (Molecules C^69^ and D^70^) were assigned to cluster #140 but have known activity with different proteins. Molecule C is a RAR*γ* antagonist of UniProt ID P13631 (retinoic acid receptor gamma).^71^ Molecule D binds to UniProt ID Q03181 (peroxisome proliferator-activated receptor delta).^72^ Cluster #140 consists of only 11 pockets from 10 unique proteins, where the predicted co-complex pockets most common for featurizing the cluster #140 pockets have PDB ligands ZHN (pentyl (3,5-dihydroxy-2-nonanoylphenyl)acetate) and VIT (vitamin e). Also in Cluster #140 are pockets on UniProt IDs P19793 (retinoic acid receptor RXR-alpha) and P48443 (retinoic acid receptor RXR-gamma). While our initial knowledge of interactions shows molecule D having activity only against Q03181, PubChem bioassay results indicate that molecule D has activity against RARA and RXRA, with potencies of 7.7 µM and 15.5 µM, respectively.^70, 72^ This is a significant finding that demonstrates a new use case, where activity not previously in our dataset was found by looking at activity for pockets in the same cluster. Both compounds were most similar to PDB ligands that were not used to characterize any of the pockets in this cluster. Molecule D was most similar to PDB ligand JNM (Tanimoto similarity 0.66); Molecule C was most similar to E9T (Tanimoto similarity 0.81). The absence of JNM and E9T as features in any of the cluster #140 pockets indicates the predictive power of the information captured by this method for pocket featurization.

As mentioned above, the AHA Protein Atlas includes docking scores and Coherent Fusion^2, 73^ machine learning scores for 6,550 compounds from the Drug Repurposing Hub. This allows quantification of how pockets in the same cluster might interact with a new compound. Using the same four-step procedure described above with regard to placing Drug-Bank compounds with their nearest cluster for all of the Drug Repurposing Hub compounds, 71520717^74^ (N-[[6-(Hydroxyamino)-6-oxohexyl]oxy]-3,5-dimethylbenzamide) as placed closest to cluster #117. Displayed in Figure 5, 71520717 has two known interactions, histone deacetylease 4 (P56524) and 5 (Q9UQL6), both of which have a pocket in cluster #117.^75^ Cluster #117 is made up of 10 total pockets from 9 different proteins and every pocket has PDB ligand ID B3N associated with its co-complex pocket matches except for Q9UQL6, which is characterized by PDB ligands SHH, 9Z8, and B3N. 71520717, like molecules A and B above, is significantly different from other ligands in the feature vector, where its best match is PDB ligand ID BHO (benzhydroxamic acid), having a Tanimoto similarity of 0.44. Among the 10 pockets which make up cluster #117, the AHA Atlas’s Coherent Fusion machine learning model and AutoDock Vina calculations highlight – with scores better than the Atlas’s threshold for more detailed binding analyses - two different proteins likely to interact with 71520717. UniProt ID Q8WUI4 (histone deacetylease 7) has the best docking score among the cluster’s pockets. PubChem’s bioassay results report a 0.17 µM “Active” result for 71520717.^76^ Activity at various levels is also shown for the other histone deacetyleases (2,6,8) in cluster #117. This evidence of cross-sensitivities in the cluster is indicative of the similarity of the pockets and the value of the information that they are grouped together. The machine learning model’s leading prediction highlights a pocket from the structural model of UniProt Q8WUI4 (metastasis-associated protein MTA3) as likely to interact with 71520717. Unlike the docking prediction, the Coherent Fusion model’s prediction does not have evidence in literature or from PubChem. Instead, assuming biological relevance of the interaction, it highlights a novel protein that might be examined for interaction.

**Figure 5:**
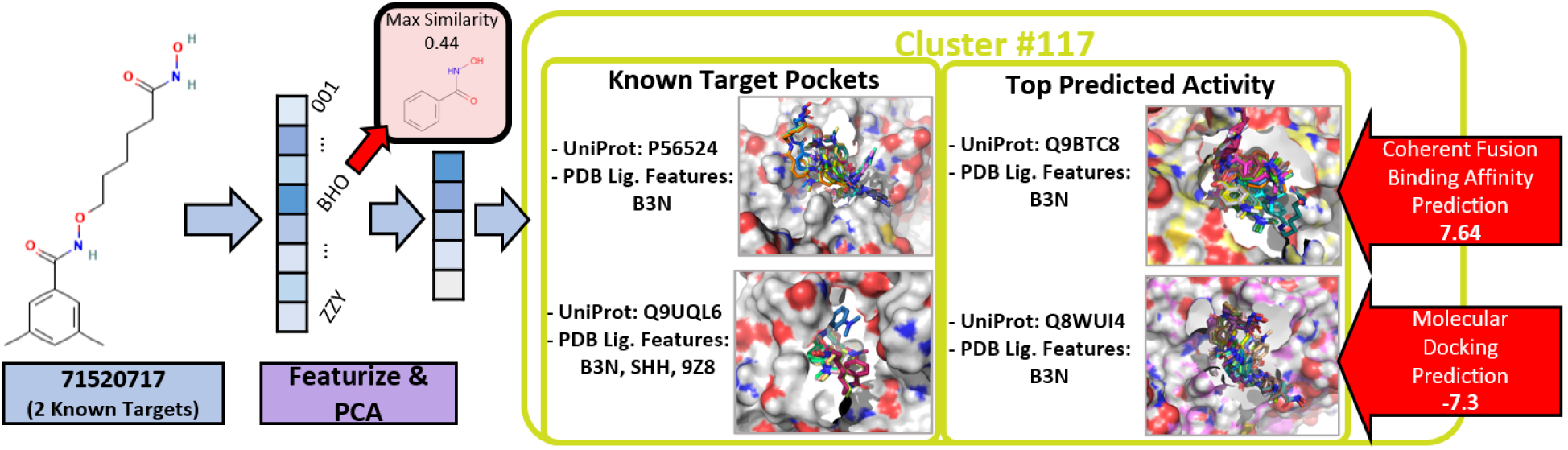
71520717 in Cluster#117 with known and top predicted activity.

## Discussion

These use cases illustrate how our ligand-based featurization and clustering approach may reveal useful information about candidate drugs and protein pocket interactions in the human proteome. While this simple approach is not perfect, it shows promise as a powerful foundational approach to improve on algorithmically. As the number of solved structures in PDB continues to grow, template-based binding site identification methods like PDBspheres will become more accurate. The 4,331 human proteins used in this work, which are provided in a supplementary file, can also be expanded beyond the AHA Atlas’s existing subset. Adding data points to this approach will serve to improve the results in Table 2. DrugBank compound results for known interactions in nearby clusters. and increase granularity. In its current state, the featurization / clustering often associates new compounds with the larger clusters. Despite controlling for cluster size in choosing the DBSCAN clustering parameters, a few clusters stand out as dominant. Cluster #1 is made up of 1,324 pockets which are characterized by template matches to NHE, HEM, 1MK, ARG, and HEA. Cluster #3 contains 653 pockets which are generally branched oligosaccharides made up of GLC, BGC, GAL, and UMQ. Finally, cluster #5 contains 1,547 pockets often associated with HEM, HEC, RLZ, JNI, and FMN. While predicting ligands to be associated with these clusters is not outright wrong, their sizes make analysis more difficult. Additional data will naturally serve to reduce this occurrence, but other clustering methods such as OPTICS^77^ may also provide an approach to keep small clusters and break up larger ones without creating unreasonable numbers of divisions. In fact, early efforts in clustering data from the AHA Atlas revealed some valuable subgroups in the larger clusters.

Nevertheless, only a subset of the possible use cases for this approach have been covered. The protein pocket clusters may have uses which span from high-throughput screening to better understanding a proteome. On the drug development side, the feature space created here can serve to aid in ligand optimization, suggest other pockets to target, and provide lead pockets for toxicity concern or assay prioritization. From a proteomic standpoint, small molecule pathways can be considered in the context of the clusters their proteins and pockets are in and vice versa. Because the feature space is traversable in terms of coordinates and distance, measurements between pockets, clusters, and regions might reveal similarities and differences in a quantifiable way. Additionally, extending this method to other proteomes might reveal nearest analogs from human to animals like non-human primates, mice, rats, etc. . . As a template-based method, recent advances in protein structure prediction will have a significant impact on our approach’s utility on proteins without known binding sites or activities.

## Conclusion

In this work, we describe and demonstrate a novel ligand-based featurization of protein pockets and clustering. The latent space captured shows strong evidence of accuracy in nominating potential multi-target candidate pockets for either designing multimodal drugs or testing in toxicity assays. The simplistic, data-driven approach developed is able to associate unseen drugs with their known target proteins and pockets, while also suggesting protein / ligand interactions which are denoted as “Active” in PubChem. The AHA Protein Atlas and binding affinity prediction methods serve to confirm validity of the clusters formed. The utility of our approach is widespread and in future work, we plan to explore additional use cases, clustering approaches, and improvements to pocket featurization.

## Data and Software Availability

The protein pockets clustered in this work come from the AHA Atlas structural models (https://doi.org/10.11578/1969730) based on the UniProt human proteome (https://www.uniprot.org/uniprotkb?query=reviewed%3Atrue%20AND%20proteome%3Aup000005640) reviewed reference set version 2020.08.19. The structural models were created using the AS2TS system (http://proteinmodel.org/). The compounds and associated protein targets used to evaluate this approach come from DrugBank version 5.1.9 (https://go.drugbank. com/releases/5-1-9) and The Drug Repurposing Hub version 3/24/2020 (https://clue. io/repurposing). The source code for PDBspheres is available at (https://github.com/ LLNL/PDBspheres). The PCA and DBSCAN clustering methods used come from scikit-learn (https://scikit-learn.org/) and the molecular docking scores were acquired via ConveyorLC (https://github.com/XiaohuaZhangLLNL/conveyorlc).

## Acknowledgement

This work was supported by American Heart Association Cooperative Research and Development Agreement TC02274. This work was performed under the auspices of the U.S. Department of Energy by Lawrence Livermore National Laboratory under Contract DE-AC52-07NA27344.

## Supporting Information Available

PDB identifiers used to filter out crystalization buffer compounds and antibody structures, PDB identifiers for the 15,489 ligands used for featurization, and a list of UniProt identifiers for proteins with a pocket successfully clustered by our method are available in the supporting information.

## Footnotes

1 1-[5-[(2,4-Difluorophenyl)thio]-4-nitro-2-thienyl]-ethanone (PubChem CID: 2819993, ^65^ Figure 4: Molecule A) and 1-(5-(2,3-Dichlorophenylthio)-4-nitrothiophen-2-yl)ethanone (PubChem CID: 46931953, ^66^ Figure 4: Molecule B), respectively.

2 Tamibarotene, 1-[1-(4-chlorophenyl)-2,5-dimethyl-1H-pyrrol-3-yl]-2-(4-hydroxypiperidin-1-yl)ethan-1-one, and 1-[1-(4-chlorophenyl)-2,5-dimethyl-1H-pyrrol-3-yl]-2-(piperidin-1-yl)ethan-1-one, respectively.

3 4-(6-Hydroxy-7-tricyclo[3.3.1.13,7]dec-1-yl-2-naphthalenyl)benzoic acid (PubChem CID: 9952709, ^69^ Figure 4: Molecule C) and 2-(4-(3-(4-Acetyl-3-hydroxy-2-propylphenoxy)propoxy)phenoxy)acetic acid (Pub-Chem CID: 6603901, ^70^ Figure 4: Molecule D), respectively.

